# Distinct and interchangeable growing patterns in colorectal cancer stem-like cells are regulated by Musashi-1

**DOI:** 10.1101/2021.11.25.469977

**Authors:** Roberto Coppo, Jumpei Kondo, Keita Iida, Mariko Okada, Kunishige Onuma, Yoshihisa Tanaka, Mayumi Kamada, Masayuki Ohue, Kenji Kawada, Kazutaka Obama, Masahiro Inoue

## Abstract

The dynamic and heterogeneous features of cancer stem-like cells (CSCs) have been widely recognized, but their nongenetic cellular plasticity mechanisms remain elusive. By using colorectal cancer organoids, we phenotypically tracked their spheroid formation and growth capacity to a single-cell resolution, and we discovered that the spheroid-forming cells exhibit a heterogeneous growth pattern, consisting of slow- and fast-growing spheroids. The isolated fast-growing spheroids seem to preserve a dual-growing pattern through multiple passages, whereas the isolated slow-growing spheroids are restricted to a slow-growing pattern. Notably, the spheroids of both patterns were tumorigenic. Moreover, the expression of CSC markers varied among the subpopulations with different growth patterns. The isolated slow-growing spheroids adopted the dual-growing pattern by various extrinsic triggers, in which Musashi-1 plays a key role. The slow-growing fraction was resistant to chemotherapy, and its successful isolation can provide an *in vitro* platform allowing us to elucidate their role in drug resistance.

## Introduction

Cancer is characterized by extensive intratumor heterogeneity (ITH) (Burrell et al., 2013; Marusyk et al., 2012; Swanton, 2012). It has been increasingly recognized that ITH contributes to drug resistance and cancer recurrence following therapy, a fact that obstructs the improvement of anticancer treatment strategies (Fisher et al., 2013; McGranahan and Swanton, 2017). ITH is known to be caused by a variety of genetic mutations, microenvironmental conditions, and cell-intrinsic plasticity (Burrell et al., 2013; Junttila and de Sauvage, 2013; Meacham and Morrison, 2013). In fact, ITH is thought to be caused by genetic mutations that have been systematically studied (Stratton et al., 2009), whereas the ITH-related nongenetic processes are currently receiving intense research attention (Brock et al., 2009; Boumahdi and de Sauvage, 2020; Marine et al., 2020; Recasens and Munoz, 2019).

Cancer stem-like cells (CSCs) are the target of one of the theories that attempts to explain the nongenetic heterogeneity of cancer (Meacham and Morrison, 2013). CSCs consist of a subpopulation of cancer cells that harbor tumor-initiating potential, extensive self-renewal capacity, and the ability to differentiate into heterogeneous cells (Hirata et al., 2019; Visvader and Lindeman, 2012). More importantly, cancer cells with stemness properties are known to be resistant to conventional therapies and responsible for tumor relapse in many cancer types (Cojoc et al., 2015; Pisco and Huang, 2015). Recently, CSC models have been revisited with the evidence that cancer cells can dynamically fluctuate from a nonstem cell-like state to a stem cell-like state (Batlle and Clevers, 2017). For this reason, the importance of the heterogeneity and plasticity within a CSC subpopulation is highlighted (Boumahdi and de Sauvage, 2020; Meacham and Morrison, 2013).

Colorectal cancer (CRC) is one of the leading causes of cancer-related death worldwide (Siegel et al., 2021). Accumulating evidence suggests that colorectal CSCs represent phenotypically dynamic (rather than static), heterogeneous cell populations that display cell plasticity characteristics (de Sousa e Melo et al., 2017; Dieter et al., 2011; Fumagalli et al., 2020; Hirata et al., 2019; Shimokawa et al., 2017). Recently, 3D cell culture systems utilizing patient-derived tumors have been developed for various cancer types, including CRC (Drost and Clevers, 2018). We, herein, use the cancer tissue-originated spheroid (CTOS) method developed by us, in which the cell–cell contact is maintained throughout the spheroid preparation, culture, and passaging (Kondo et al., 2011). The growth of each CRC CTOS within the same line is quite heterogeneous (Kondo et al., 2019), suggesting that CRC CTOSs can retain heterogeneous populations of cells. We have also demonstrated that a small subset of cells within the CRC CTOSs can initiate regrowth after exposure to high-dose radiation, an observation that was nongenetically and reversibly determined (Endo et al., 2020).

As current cancer therapies have been designed and developed mainly against the fast-growing cancer cells, the importance of the quiescent or slow-growing CSCs has been overlooked. Consequently, these slow-growing CSCs may survive the anticancer treatment, revert to the fast-growing cancer cells, and serve as a reservoir for tumor regrowth (Barriga et al., 2017; de Sousa e Melo et al., 2017; Kurtova et al., 2015; Rehman et al., 2021; Roesch et al., 2010). To characterize the dynamics of quiescent or slow-growing CSCs, an analysis at the single-cell resolution is required because the nature of the slow-growing cells is usually masked by that of the fast-growing cells. Recently, single-cell transcriptome analyses were recruited and served the study of the characteristics of the slow-growing CSCs (Shiokawa et al., 2020). However, such a “snapshot” analysis involving a procedure for isolating CSCs with the use of markers (Shiokawa et al., 2020) or dye retention (Dembinski and Krauss, 2009; Richichi et al., 2013) exerts limitations when applied to a continuously changing process. Therefore, the employment of a phenotypically trackable cell culture system allowing for a single-cell resolution is necessary.

Spheroid formation from a single cell is one of the phenotypical hallmarks of CSCs (Pastrana et al., 2011; Todaro et al., 2007; Vermeulen et al., 2008). In this study, by using CRC organoids, we could (i) track the capacities of both the spheroid formation and growth with a single-cell resolution and (ii) reveal the existence of heterogeneous subpopulations among the CSCs, consisting of fast-and slow-growing cells. The cells derived from the fast-growing spheroids gave rise to both slow- and fast-growing spheroids, by adopting a “dual-growing pattern.” Meanwhile, the cells from the slow-growing spheroids gave rise to slow-growing spheroids only, thus following a “slow-growing pattern.” Once adopted, the slow-growing pattern seemed to be preserved over the spheroid passages. We could demonstrate the phenotypic differences between the growth patterns as well as their interchangeable features when following these patterns, in which Musashi-1 (MSI1) was identified as playing a functional role. Moreover, the successful cell culturing of the slow-growing fraction of CSCs enabled us to investigate the plasticity of CSCs; the latter could play a critical role in drug resistance and tumor recurrence.

## Results

### Heterogeneous growth ability of CRC cells at a single-cell level

To precisely track the growth ability of CRC cells in CTOSs at a single-cell resolution, we modified the conventional spheroid-forming assay and developed a single-cell-derived spheroid-forming and growth (SSFG) assay, which includes the undertaking of (i) a mild dissociation of the CTOS, (ii) an initial confirmation of the single-cell status within a well, (iii) the establishment of growth permissive culture conditions, and (iv) a time-course analysis of each well (Figure 1A). We applied the SSFG assay to a CRC CTOS line, C45, which in turn demonstrated a wide range of CTOS growth abilities (Kondo et al., 2019). The SSFG assay enabled us to exclude nonsingle cells at the very beginning of the assay (Figure 1B), as well as the nongrowing cells (Figures 1C and 1D) that presented with several patterns: early death (Figure S1A-a), late death (Figure S1A-b), growth arrest (Figure S1A-c), and a decline in size (Figure S1A-d). The maximum area of the nongrowing spheroids was below 2.5 × 10^3^ μm^2^. The spheroid-forming capacity–a widely accepted feature of stem cells–was, on average, 59%. The single-cell-derived growing spheroids of the C45 CTOSs exhibited a substantial range of growth (Figures 1C–1F), with a 228-fold width. We could measure the growth variation of single cells through the SSFG assay in 13 additional lines of CRC CTOSs from different patient tumors (Table S1, Figures 1G–1I, and S1B–S1K). The spheroid-forming capacities were between 19% and 59% (Table S2). The maximum growth ability varied among the studied lines. In all 14 studied CRC CTOS lines (including C45), the sizes of the spheroids within each line varied substantially, and two of the lines (namely, C120 and C132) demonstrated a statistically significant bimodal distribution. The mutational profile (Table S3) showed no clear correlation with the spheroid-forming capacity, the growth, or the growth range. These results indicate that the spheroid-forming cells (SFCs) in the CRC CTOSs consist of cells with different growth abilities.

**Figure 1.**
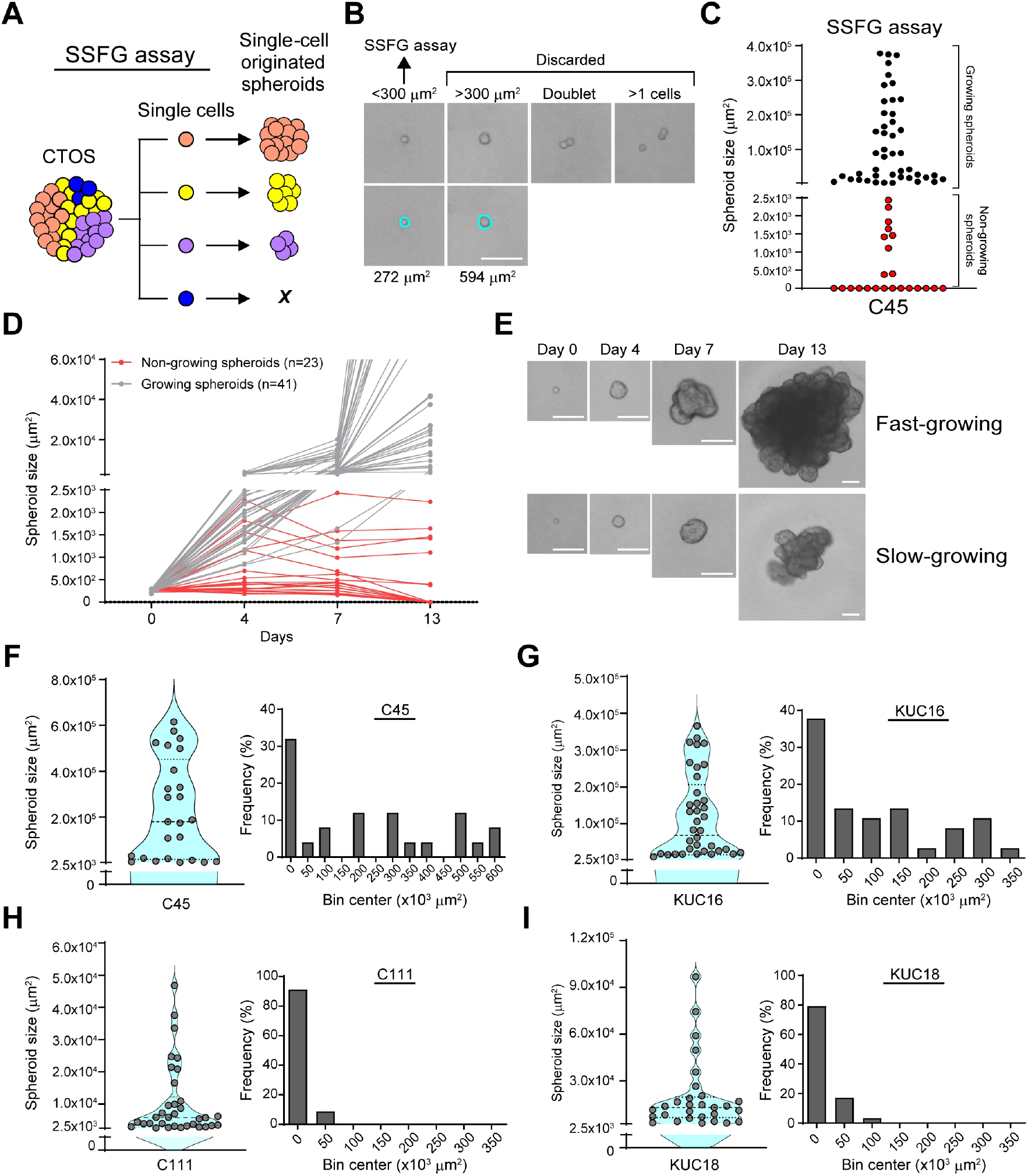
CRC organoids exhibit heterogeneous growth at the single-cell level. (**A**) Schematic overview of the SSFG assay undertaken. A circle represents a cell; different colors represent heterogeneous populations of the cells. (**B**) Phase-contrast images of the SSFG assay at day 0. The blue circles indicate the surface area of single cells. (**C**) Size of the C45 single-cell-derived spheroids in the SSFG assay. Gray and red dots represent growing and nongrowing spheroids, respectively. (**D**) Growth curve of the growing (gray) and the nongrowing (red) spheroids from (1C). (**E**) Phase-contrast images of the growing spheroids in (1C). (**F–I**) Left panel: a violin plot of the SSFG assay. Right panel: frequency distribution of the size of the growing spheroids. All scale bars in Figure 1: 100 μm.

### SFCs consisted of two subgroups with different growth phenotypes

In an attempt to investigate cell growth heterogeneity in more detail, we selected several individual C45 spheroids at the end of the SSFG assay. As each spheroid was strictly derived from a single cell, it can be called a “clone.” We selected two clones (C45-1 and C45-2) from the relatively slow-growing C45 spheroids and two clones (C45-3 and C45-4) from the fast-growing spheroids (Figure 2A). The growth was clearly different among the studied groups (Figures 2B and S2A), with a similar appearance (Figures 2B and S2B). We expanded each clone *in vitro* and performed a second round of SSFG assays. The spheroid-forming capacity was preserved in all clones (Figure 2C). The slow-growing spheroids gave rise to only slow-growing spheroids (slow-growing pattern, S-pattern) (Figure 2D). Surprisingly, the fast-growing spheroids gave rise to both slow- and fast-growing spheroids (dual-growing pattern, D-pattern) (Figure 2D). We then set the putative threshold between the two phenotypes at 1.0 × 10^5^ μm^2^, based on the rounded value of the maximum size of the C45-1 spheroids at day 13. We could confirm the D-pattern in three other CRC CTOS lines (Figures 2E, 2F, and S2C-2F). To further investigate the stability of the aforementioned growth features, we performed additional rounds of the SSFG assay on the C45. In all four C45 clones, the growth pattern was preserved during all three rounds of the SSFG assay (Figures 2G–2J). These results indicate that the SFCs within each CRC CTOS consisted of two different subgroups each expressing the S- and the D-pattern, thus demonstrating stem cell heterogeneity and plasticity features.

**Figure 2.**
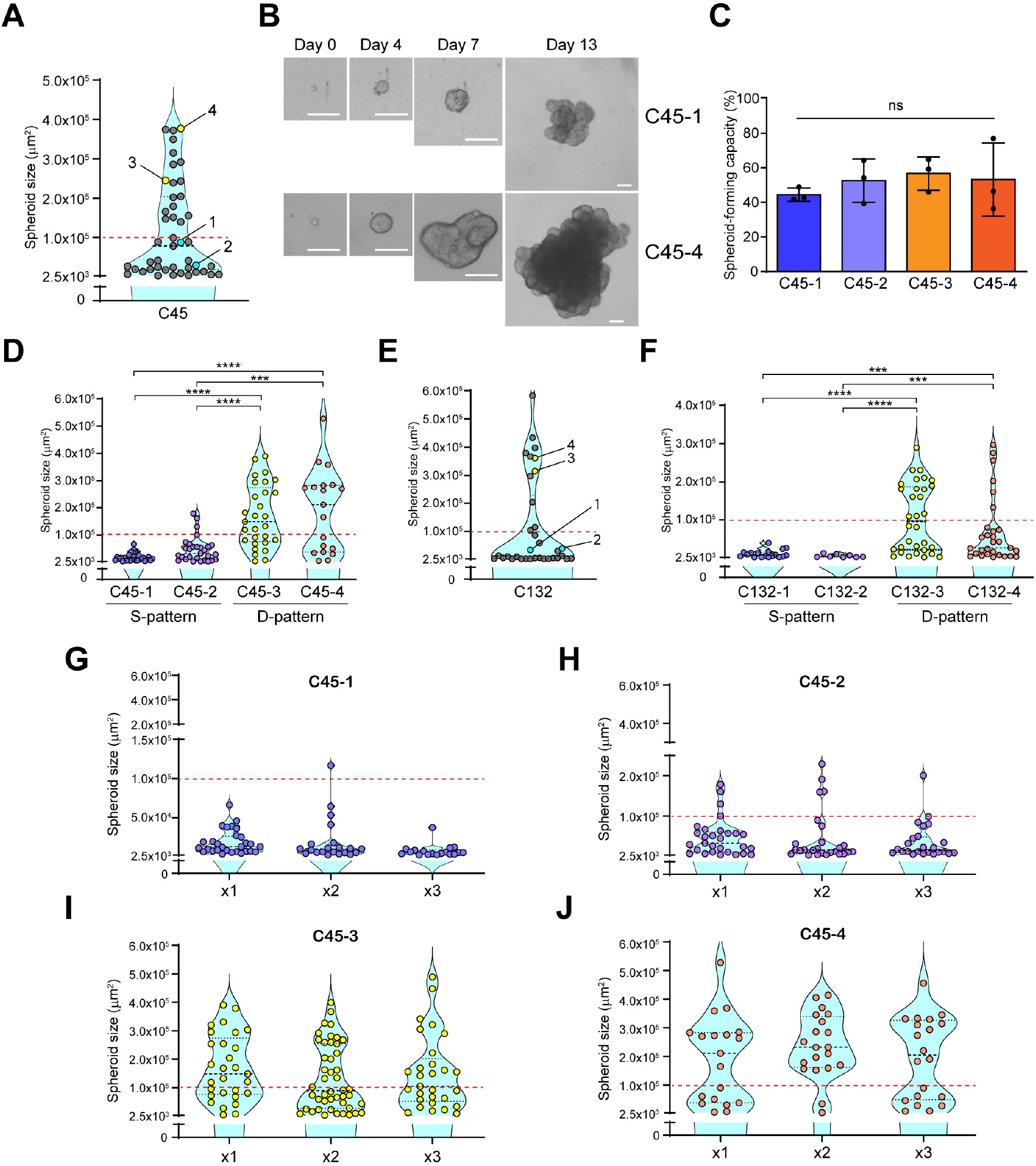
CRC spheroids demonstrate different but stable growth capacities. (**A**) Violin plots of the SSFG assay for the C45 CRC CTOS line. Four selected spheroids are indicated. The red dashed line indicates a rounded value (1.0 × 10^5^ μm^2^) of the area at day 13, for the C45-1 line, and it is indicative of a putative threshold between the slow- and the fast-growing spheroids in the C45 lines. (**B**) Time-course images of the C45-1 and C45-4 lines. Scale bars: 100 μm. (**C**) Spheroid-forming capacity of the indicated clones. The mean ± SD is shown, tested by one-way analysis of variance (ANOVA), followed by Tukey’s test. (**D**) Violin plots of the SSFG assay for the indicated clones. (**E**) Violin plots of the SSFG assay for the C132 line. Four selected clones are highlighted. (**F**) Violin plots of the SSFG assay for the indicated clones. (**G–J**) Violin plots of the serial SSFG assays for the indicated clones. The first round is indicated as “x1,” the second as “x2,” and the third as “x3.” The results of the first round are the same as in (2D).

### Slow-growing spheroids retained a potential to transition to the dual-growing pattern

Subsequently, we investigated the transition between the S- and the D-patterns. We used C45-4 spheroids (which exhibited a D-pattern) and subjected them to another round of the SSFG assay, from which we isolated slow-(C45-4S) and fast-growing (C45-4F) spheroids (Figure 3A). We then repeated the selection three times (Figure 3A). During the multiple rounds of the SSFG assay, the D-pattern was preserved in the C45-4F spheroids. On the other hand, the C45-4S spheroids exhibited the D-pattern in the first round, then gradually lost the fast-growing spheroids, and reached growth plateau by adopting the S-pattern in the third round (see the control groups in the following results of the SSFG assay using passaged C45-4S spheroids). After three rounds of the SSFG assay, we named the subgroups derived from the C45-4S and the C45-4F spheroids as “C45-4SR” (slow-growing restricted) and “C45-4D” (dual-growing) spheroids, respectively. The spheroid-forming capacity of both subgroups did not differ from that of the parent C45-4 (see Figure 2C and the control groups in the following results of the SSFG assay). As the C45-4 spheroids were derived from a single clone, we assumed that the growth pattern is likely to be regulated by nongenetic mechanisms.

**Figure 3.**
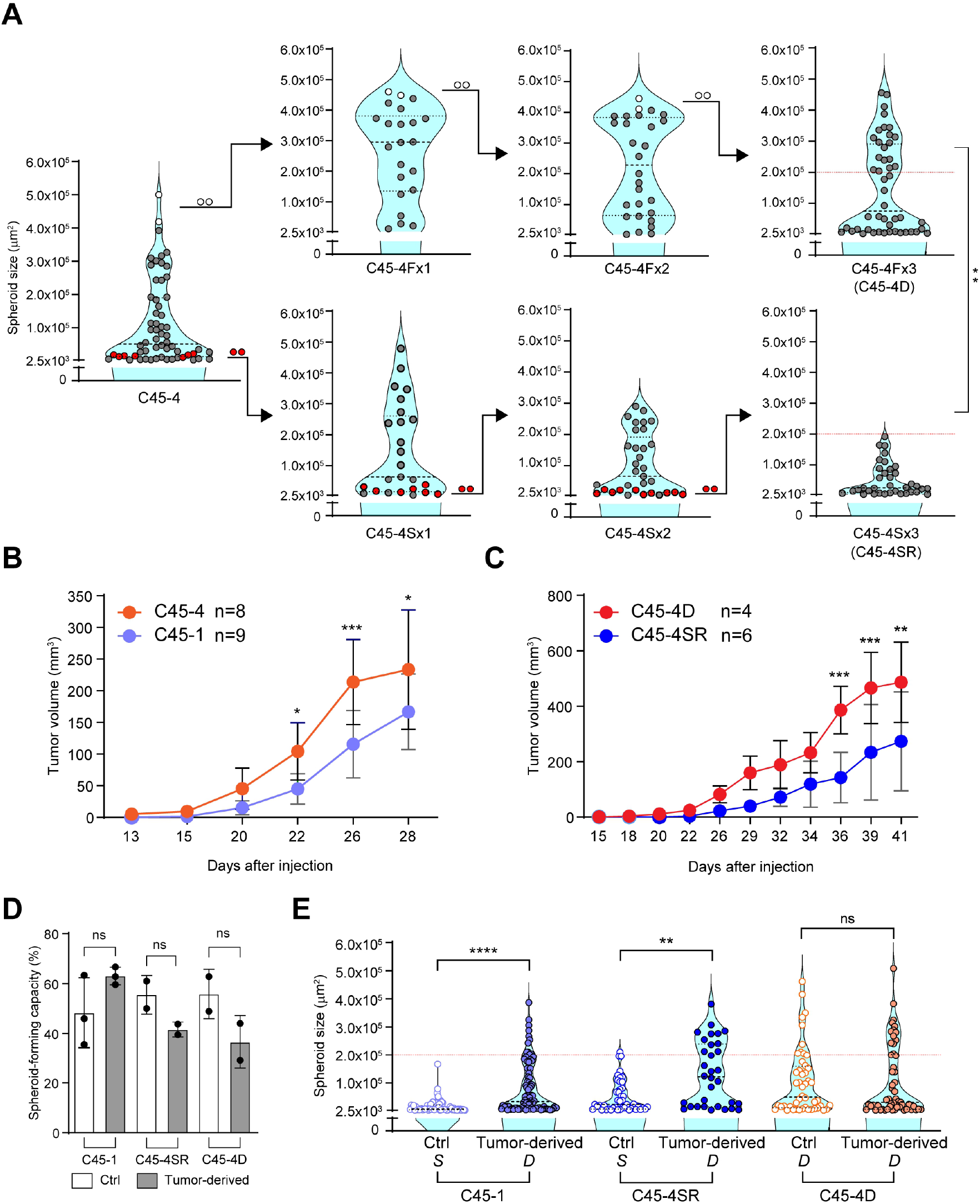
CRC spheroids show cell plasticity in their growth characteristics. (**A**) Violin plots of the SSFG assay for the indicated clones. Red and open circles represent the selected slow-growing (C45-4S) and fast-growing (C45-4F) spheroids, respectively. Several fast- and slow-growing spheroids with similar size were collected separately and subjected to the next round of the SSFG assay. Three rounds of the SSFG assay were performed for each subclone. The first round is indicated as “x1” (C45-4Fx1, -4Sx1), the second as “x2” (C45-4Fx2, -4Sx2), and the third as “x3” (C45-4Fx3, -4Sx3). The red dotted line expresses the rounded value (2.0 × 10^5^ μm^2^) of the area at day 13, for the C45-4Sx3 line, and it is indicative of a putative threshold between the slow- and fast-growing spheroids in the C45-4 lines. (**B**, **C**) Growth curves of xenograft tumors originating from C45-1 and -4 (3B) and C45-4D and -4SR (3C) spheroids. The mean ± SD is shown; n, the number of animals in each group; statistical analyses were performed by two-way ANOVA, followed by Bonferroni’s test. (**D, E**) Spheroid-forming capacity (3D) and violin plots of the SSFG assay (3E) for the C45-1, C45-4SR, and C45-4D subclones comparing control CTOSs maintained *in vitro* (Ctrl) with those prepared from the xenografts in (3B, 3C) (tumor-derived). *S*, S-pattern; *D*, D-pattern.

We next examined the tumorigenicity of the subclones. To do that, we subcutaneously injected the spheroids of C45-1 and C45-4SR from the S-pattern group and C45-4 and C45-4D from the D-pattern group into immunodeficient mice. All spheroids were tumorigenic (Figures 3B and 3C). Although the S-pattern group demonstrated growth retardation compared with the D-pattern group, the group’s growth rate eventually caught up. We then prepared CTOSs from the xenografts and subjected them to the SSFG assay. The spheroid-forming capacity was preserved after the xenograft formation (Figure 3D). The D-pattern was preserved in the C45-4D spheroids (Figure 3E). Interestingly, the slow-growing C45-1 and C45-4SR spheroids acquired the D-pattern phenotype following the xenograft formation (Figure 3E). These results indicate that both spheroids with the S- and D-patterns were tumorigenic. More importantly, the S-pattern groups adopted a D-pattern after interacting with the tumor microenvironment, thereby proving their stemness plasticity.

### Transition between growth patterns was regulated by cell–cell contact through Notch signaling

As the S-pattern was maintained following the spheroid isolation, we speculated that the transition mechanisms from the S- to the D-patterns might involve a cell–cell interaction. We, thus, generated chimeric spheroids by aggregating the EGFP-labeled C45-4SR cells and the mCherry-labeled C45-4D cells (Figures 4A and 4B), and subjected them to the SSFG assay. The EGFP-labeled C45-4SR cells acquired the D-pattern phenotype within the same chimeric spheroids, in the same way as the mCherry-labeled C45-4D cells (Figures 4C and 4D). On the other hand, the S-pattern was maintained in the EGFP-labeled C45-4SR cells when both types of cells were co-cultured in different gel droplets within the same wells (Figures 4E and 4F), thus indicating that a direct cell–cell interaction was necessary for this transition. To further investigate the molecular mechanisms of this cell–cell interaction, we proceeded to examine the Notch signaling, which plays an important role in determining the cell fate (Bu et al., 2013; Srinivasan et al., 2016a; Srinivasan et al., 2016b). The C45-4D and C45-SR spheroids were treated overnight with a Notch inhibitor, DAPT (50 μM), and then generated chimeric spheroids. The inhibition of the Notch signaling by DAPT was confirmed through the Western blotting of the Notch intracellular domain (NICD) (Figure 4G). Although the *in vitro* treatment with DAPT did not affect the spheroid-forming capacity (Figure 4H), it managed to inhibit the transition of the C45-SR cells to the D-pattern within the chimeric spheroids (Figure 4I). Thus, the Notch signaling was necessary for the switching of the SFCs from the S- to the D-pattern; by doing so, it also provided us with further evidence of the SFC stemness plasticity.

**Figure 4.**
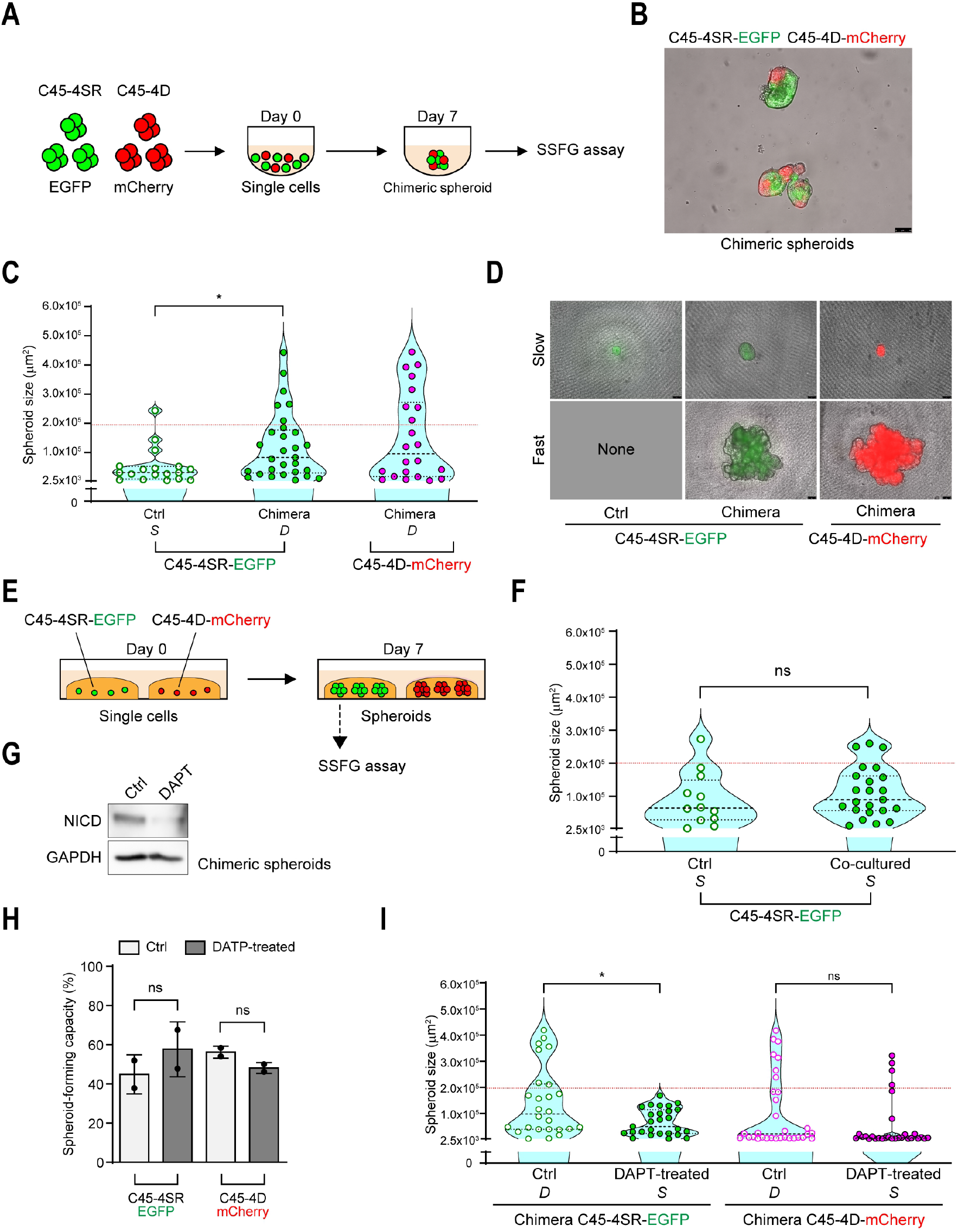
The Notch pathway is necessary for the cell growth plasticity of CRC spheroids induced by cell–cell interactions. (**A**) Schematic overview of the chimeric spheroid experiments that enabled physical cell–cell interactions between cells of different nature; mCherry-labeled C45-4D (red circles) and EGFP-labeled C45-4SR (green circles) cells. (**B**) Representative image of the C45-4D-mCherry/C45-4SR-EGFP chimeric spheroids. Scale bar: 75 μm. (**C**) Violin plots of the SSFG assay comparing the cells from pure C45-4SR-EGFP (Ctrl), the EGFP-positive cells from the C45-4D-mCherry/C45-4SR-EGFP chimera (Chimera), and the mCherry-positive cells from the chimera. (**D**) Images of the slow- and fast-growing spheroids, at day 13, derived from EGFP- or mCherry-positive cells. (**E**) Schematic overview of the co-culture system without cell–cell interactions; mCherry-labeled C45-4D cells, red; EGFP-labeled C45-4SR cells, green. (**F**) Violin plots of the SSFG assay comparing the EGFP-positive cells from the pure C45-4SR-EGFP spheroids (Ctrl) with those from the co-cultured spheroids. (**G**) Western blot analysis of the expression of NICD in C45-4D-mCherry/C45-4SR-EGFP chimeric spheroids treated with DMSO (0.1%) or DAPT (50 µM) for 7 days. (**H, I**) Spheroid-forming capacity (4H) and violin plots of the SSFG assay (4I) comparing the DMSO-treated EGFP-positive cells with the DAPT-treated C45-4D-mCherry/C45-4SR-EGFP chimeric spheroids (Chimera), and the DMSO-treated mCherry-positive cells with the DAPT-treated chimeric cells. *S*, S-pattern; *D*, D-pattern.

### MAPK/ERK is necessary for growth of fast-growing spheroids but not of the slow-growing ones

We examined the intracellular signaling–the MAPK/ERK and AKT/mTOR pathways–in the aforementioned subgroups. Among the signaling molecules, the levels of protein and the phosphorylation of MEK1/2 and ERK1/2 were found to be lower in the slow-growing cells (Figure 5A). To examine the role of the MEK/ERK pathway in the transition of the growth patterns, we introduced an MEK1/2 inhibitor, PD0325901 (MEKi), in the SSFG assay. In some cases, the spheroid-forming capacity decreased slightly as a result of the MEKi treatment (Figure 5B), and no morphological changes were observed (Figure 5C). The results of the SSFG assay in the C45-1 spheroids did not change, even after their incubation with 100 nM of MEKi (Figure 5D). In the C45-4SR (Figure 5E) and the C45-4D (Figure 5F) spheroids, the relatively fast-growing spheroids decreased along with the increase of the *in vitro* MEKi concentration; however, the slow-growing spheroids remained unaffected. The experimental design and the results of these experiments are summarized in Figure S3A. Other D-pattern lines (such as the parental C45, C132, and KUC16 lines) responded to the MEKi in a manner similar to the C45-4D spheroids: the fast-growing spheroids decreased in the SSFG assay (Figure S3B), without demonstrating any differences with regard to their spheroid-forming capacity (Figure S3C) or their morphology (Figure S3D).

**Figure 5.**
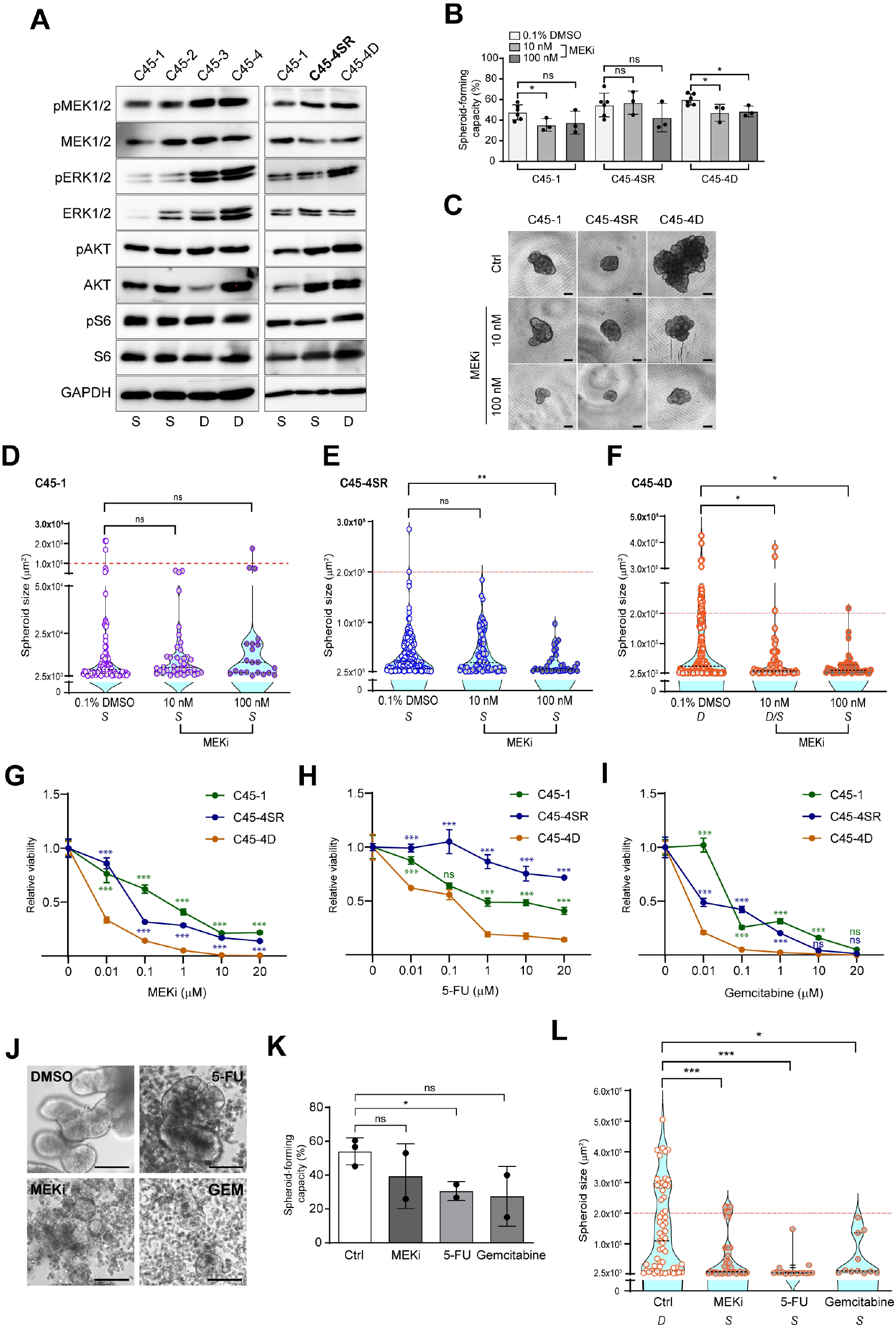
Slow-growing CRC spheroids show resistance to an MEK1/2 inhibitor with a single-cell resolution. (**A**) Western blots analysis of the intracellular signaling in the spheroids of the indicated clones after 7 days of culture. S, slow-growing; D, dual-growing. (**B**) Spheroid-forming capacity of the C45-1, C45-4SR, and C45-4D clones treated with DMSO (0.1%) or MEKi (10 and 100 nM) for 13 days. Schematic overview of the experimental design, and the results are presented in Figure S3A. (**C**) Representative phase-contrast images of the spheroids derived from single cells of the indicated lines at day 13, treated as indicated. Scale bars: 100 μm. (**D–F**) Violin plots for C45-1 (5D), C45-4SR (5E), and C45-4D (5F), comparing the DMSO (0.1%) treatment with the MEKi (10 or 100 nM) treatment; all treatments lasted for 13 days. Representative data of three independent experiments are shown. The red dashed line in (5D) is the putative threshold for the C45-1 line, whereas those lines in (5E, 5F) are the putative thresholds for the C45-4SR and C45-4D lines. *S*, S-pattern; *D*, D-pattern; D/S, intermediate pattern. (**G–I**) Dose-dependent curves of the indicated C45 subclones treated with MEKi (**G**), 5-FU (**H**), and gemcitabine (**I**), as evaluated by an ATP assay. The mean ± SD is shown, tested by two-way ANOVA, followed by Tukey’s test. Comparisons of C45-1 *vs.* C45-D and C45-4SR *vs.* C45-D are shown. (**J**) Representative phase-contrast images of the C45-4D spheroids (at day 7), treated with the indicated drugs. Scale bars: 100 μm. (**K, L**) Spheroid-forming capacity (5K) and violin plot of the SSFG assay (at 13 days) (5L) for the C45-4D subclone, comparing the DMSO (0.1%)-treated with indicated drug-treated spheroids from Figure 5J. *S*, S-pattern; *D*, D-pattern.

### Slow-growing spheroids contained more drug-resistant cells in CTOS

Subsequently, we assessed the subpopulations for different growth characteristics (namely, C45-1, C45-4SR and C45-4D) with the conventional sensitivity assay of CTOSs (Kondo et al., 2019). We then tested the MEKi and two chemo-drugs: 5-FU (a drug that is currently used in clinical practice) and gemcitabine (an effective drug toward C45 but still well tolerated–even at high doses–by a substantial fraction of cells, as shown in our previous study) (Kondo et al., 2019). The S-pattern spheroids (C45-1 and C45-4SR) were significantly more resistant to these drugs than the D-pattern spheroids (C45-4D) (Figures 5G–5I). Notably, at higher doses of each drug, a number of small intact spheroids remained in the C45-1, the C45-SR, and even the C45-4D spheroids (Figure 5J). We then performed SSFG assays for the remaining small spheroids of C45-4D after exposure to each drug. In some cases, the spheroid-forming capacity was decreased slightly with the drug treatment (Figure 5K). The SSFG assay revealed that the remaining small spheroids consisted only of the slow-growing cells (Figure 5L). These results suggest that the slow-growing spheroids contain more drug-resistant cells, and that the sensitive fraction in the D-pattern cells consisted of fast-growing spheroids.

### Transcriptome profiling revealed a heterogeneous expression of CSC marker genes, and highlighted MSI1 as a candidate molecule for regulating the growth pattern transition

In an attempt to shed more light on the molecular characteristics of the subclones with the distinct growth features, we analyzed the differentially expressed genes between the subgroups. The single cells derived from the C45-1, C45-4SR, and C45-4D subclones were cultured for 7 days and were subsequently subjected to microarray analyses. The volcano plot analyses revealed the similarity of their gene expression profiles and the existence of some differentially expressed genes (Figures 6A–6C). Of the 29,596 genes examined, 408 genes were found to be upregulated more than 1.5-fold, and 620 genes were found to be downregulated less than 0.67-fold when the three aforementioned subclones were compared (Figure 6D, Table S4). Surprisingly, among these differentially expressed genes, many have been previously reported as stem cell markers of CRC (Batlle and Clevers, 2017; Dalerba et al., 2011; Hirata et al., 2019; O’Brien et al., 2007; Ricci-Vitiani et al., 2007). The levels of MSI1, MEX3A, SOX4, EPHA4, and LRIG1 were higher in the C45-4D spheroids; the level of PTPPRO was higher in the C45-1 spheroids; and the levels of LGR5, PROM1 (CD133), and RGMB were higher in the C45-4SR spheroids (Figures 6A–6C). We could confirm the expression patterns of LGR5, PROM1, and MSI1 by undertaking a semiquantitative RT-PCR (Figure 6E). The results suggested that the expression levels of the reported CSC genes changed according to the status of the SFCs. By testing the hallmark collection in MSigDB v7.4 (https://www.gsea-msigdb.org/gsea/msigdb/index.jsp), we could conclude that the MYC signature (Zeller et al., 2003) was significantly enriched in the C45-4D spheroids compared with the C45-1 or the C45-4SR spheroids (Figure 6F). Indeed, the protein levels of MYC were higher in the C45-4D spheroids than in the C45-4SR spheroids, and very low in the C45-1 spheroids (Figure 6G); the latter seems to be reasonable as MYC lies at the crossroads of many growth-promoting signal transduction pathways (Dang, 2012). To further explore the genes that characterize the different phenotypes of SFC growth, we selected those probe sets with relatively high intensity. Following a clustering and a heatmap analyses of the 148 selected genes (Table S5), we successfully identified the selection of the differentially expressed genes, in which *MSI1* and *MEX3A* were included (Figure 6H).

**Figure 6.**
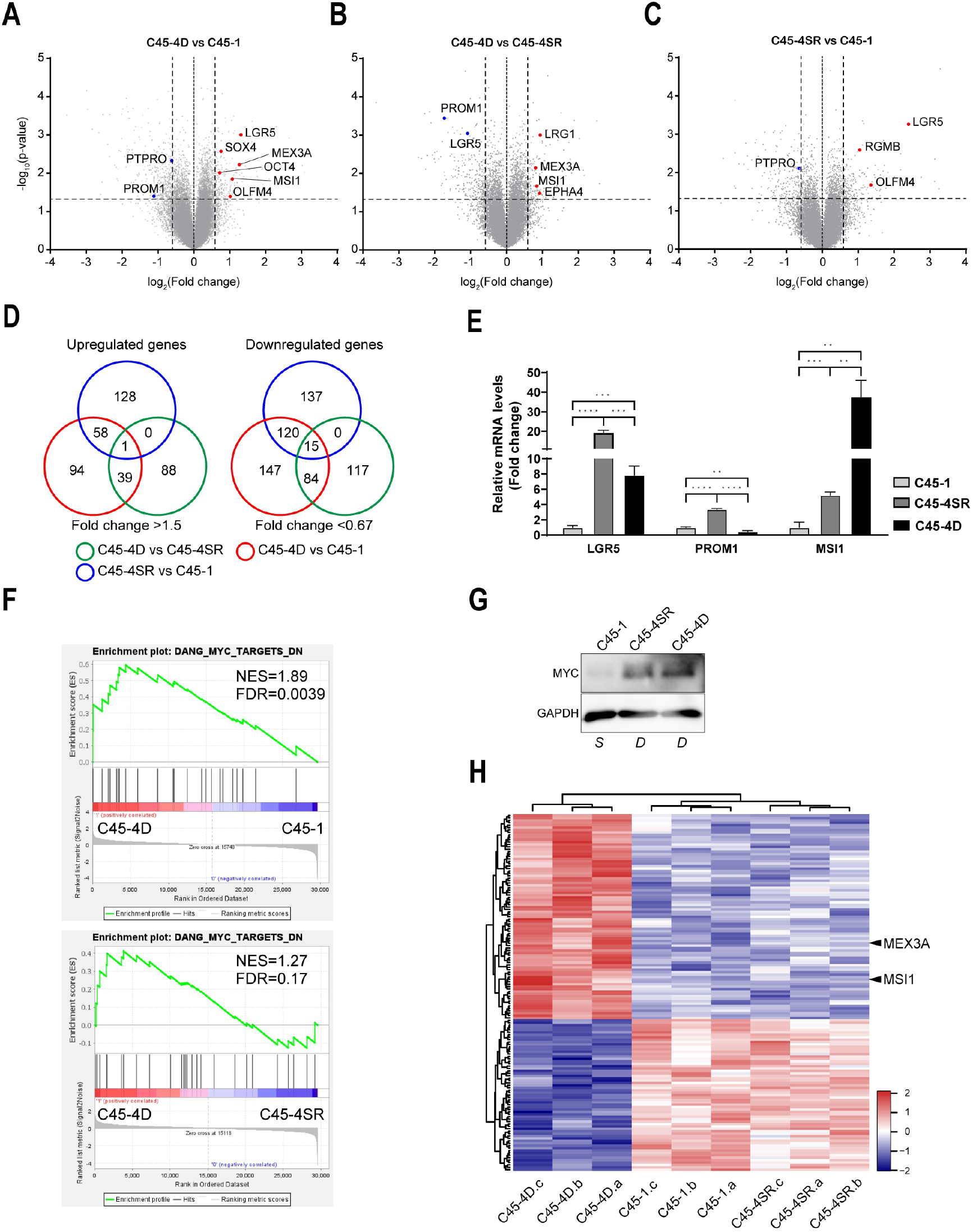
Transcriptome analysis revealed a heterogeneous expression of stemness-related genes among the CRC spheroids. (**A–C)** Volcano plots displaying the differentially expressed genes from gene expression microarray data obtained from three independent experiments (n = 3). (6A) C45-4D *vs.* C45-1, (6B) C45-4D *vs.* C45-4SR, and (6C) C45-4SR *vs.* C45-1. The red and blue dots represent the upregulated and downregulated colorectal CSC marker genes, respectively. The threshold lines are indicated as ± 0.59 for the log2 (fold change) and as 1.3 for the − log10 (p-value) **(D)** Venn diagram presenting the number of significantly upregulated (left panel) and downregulated (right panel) genes in the comparison of C45-1, C45-4SR, and C45-4D spheroids. (**E**) Relative mRNA levels of the LGR5, PROM1 (CD133), and MSI1 genes in the indicated C45 clones after 7 days of culturing. (**F**) Gene set enrichment analysis (GSEA) of the transcriptome data by using the c-Myc target gene signature (Zeller et al., 2003; DANG_MYC_TARGETS_DN), and by comparing the dual-growing C45-4D with the slow-growing C45-1 (upper panel) or C45-4SR (lower panel). (**G**) Western blot analyses of the c-MYC protein levels in C45-1, C45-4SR, and C45-4D spheroids after 7 days of culturing. S, S-pattern spheroids; D, D-pattern spheroids. (**H**) Heatmap and clustering analyses. The probe sets were selected with a relatively high intensity: top 40% of mean expression values, and FDR < 0.1 in the comparison of C43-4D with both the C45-1 and C45-4SR lines. MSI1 and MEX3A are indicated.

### MSI1 was a key factor in the transition between the different growth patterns

The posttranscription regulator MSI1 is an RNA-binding protein that is reportedly a CSC marker in the normal intestine and the CRC (Nishimura et al., 2003; Potten et al., 2003; Schulenburg et al., 2007). MSI1 is considered to be involved in the homeostasis of the intestine as a regulator of the self-renewal of stem cells in both the normal intestine (Yousefi et al., 2016) and during the growth of a CRC (Li et al., 2015; Sureban et al., 2008). The expression levels of MSI1 were confirmed to be higher in the C45-4D spheroids than in the C45-1 and C45-4SR spheroids (Figure 7A). This was also true in the subclones of other lines, such as C132 and the KUC16 (Figure 7B). The phosphorylation of ERK1/2 and the protein levels of MYC were increased in parallel to the MSI1 levels (Figure 7B). We assessed the change in the MSI1 protein levels through MEK1/2 inhibition. When MEK1/2 was inhibited at a lower concentration, the MSI1 levels increased along with the MYC decrease; however, at a higher concentration, the protein levels of MSI1 and MYC both decreased in the C45-4D spheroids (Figure 7C). To further investigate the functional role of MSI1 in the transition between the different growth patterns, we knocked out the MSI1 gene in the C45-4D spheroids by using the CRISPR/Cas9 system, generating the C45-4D_sgMSI1 spheroids. The *MSI1* knockout in C45-4D resulted in an increase of the occurring ERK1/2 phosphorylation and a decrease in the MYC protein levels (Figure 7D). No difference was observed with regard to the spheroid-forming capacity (Figure 7E), whereas the ability to generate fast-growing spheroids was impaired (Figure 7F). Notably, the spheroid-forming capacity was unaffected, thus suggesting that MSI1 is involved in the transition of the growth pattern but not in the capability of spheroid formation. Subsequently, we examined whether the cell–cell contact can rescue the impaired transition of the C45-4D_sgMSI1 cells to the D-pattern, as shown in Figure 4C. For this reason, we generated chimeric spheroids by mixing tRFP-labeled C45-4D_sgMSI1 cells and wild-type C45-4D cells, and subjected them to the SSFG assay. The C45-4D_sgMSI1 cells could not demonstrate their D-pattern phenotype, even in the presence of wild-type cells (Figure 7G), thereby suggesting that MSI1 is involved in the cell–cell contact-induced transition from the S- to the D-pattern of growth. We then overexpressed MSI1 in slow-growing C45-4SR and generated the C45-4SR_MSI1OE spheroids, which we similarly subjected to the SSFG assay. MSI1 overexpression in the C45-4SR spheroids resulted in an increase of the ERK1/2 phosphorylation and the MYC protein levels (Figure 7H). No difference was observed with regard to their spheroid-forming capacity (Figure 7I), whereas the fast-growing spheroids were increased in the C45-4SR_MSI1OE cultures (Figure 7J). Notably, the C45-4SR_MSI1OE spheroids demonstrated the D-pattern growth phenotype, but not all their cells gave rise to fast-growing spheroids. This observation suggested that MSI1 is involved in the transition of the growth pattern and not simply in the growth of the spheroids. Taken together, MSI1, ERK1/2, and MYC seem to be tightly linked, although they are likely to be regulated by complex networks. Moreover, MSI1 clearly plays a functional role in the transition between the different growth patterns in CRC CTOSs.

**Figure 7.**
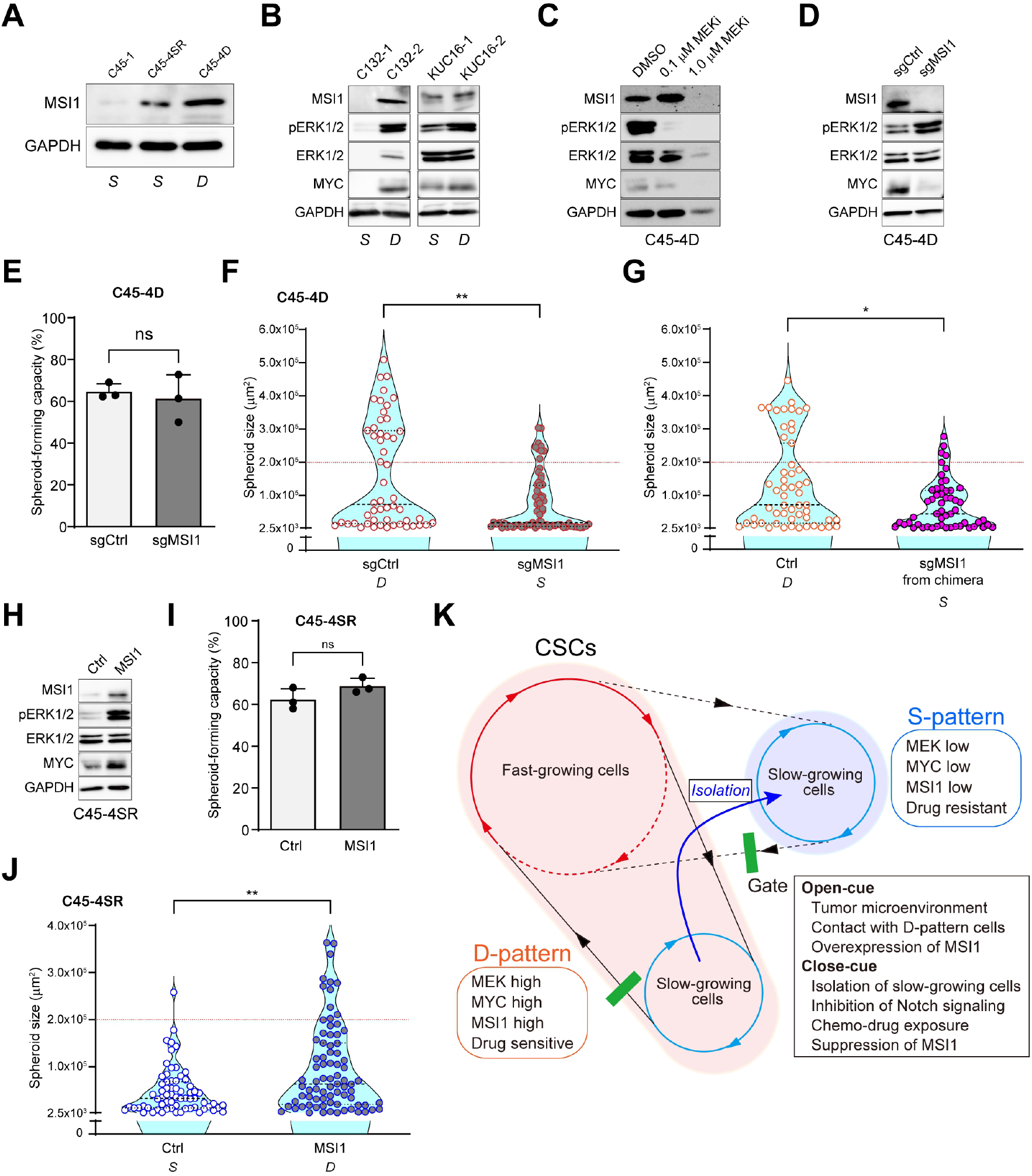
The RNA-binding protein MSI1 regulates the cell growth and plasticity of CRC spheroids. (**A**, **B**) Western blot analyses of the MSI1 protein from C45 spheroids in slow- (S) and dual- (D) growing pattern. (**B**) Western blot analyses of the indicated proteins in other lines after 7 days of culturing. (**C**) Western blot analyses of indicated proteins in the C45-4D subclone. C45-4D spheroids were incubated with 0.1% of DMSO, and 0.1 or 1.0 µM of MEKi, for 7 days. (**D**) Western blot analyses of MSI1 and indicated protein levels in C45-4D spheroids infected with lentiviruses expressing Cas9 and the MSI1 sgRNA (sgMSI1) or nontargeting sgRNA (sgCtrl). (**E**, **F**) Spheroid-forming capacity (7E) and violin plot of the SSFG assay (7F) for the C45-4SR subclones comparing the control and the *MSI1* knockout cells. (**G**) Violin plot of the SSFG assay for the C45-4D tRFP-labeled sgRNA MSI1 (sgMSI1) cells derived from the chimeric spheroids mixed with C45-4D wild-type cells (Ctrl). (**H**) Western blot analyses of MSI1 and other indicated proteins in C45-4SR spheroids infected with a lentivirus that constitutively expresses MSI1 or with the corresponding empty vector (Ctrl). (**I**, **J**) Spheroid-forming capacity (7I) and violin plot of the SSFG assay (7J) for the C45-4SR subclones, comparing the control and the MSI1 overexpressing cells. *S*, S-pattern; *D*, D-pattern. (**K**) Schematic model of the dual mode of growth in CRC CSCs.

## Discussion

Our experiments revealed that the patient-derived CRC organoids consist of phenotypically heterogeneous and interchangeable SFCs with different growth capacities. The slow-growing cells can give rise only to slow-growing restricted spheroids (S-pattern), whereas the fast-growing cells can give rise to both fast- and slow-growing spheroids (D-pattern). The transition from the S- to the D-pattern is molecularly regulated, and MSI1 seems to play a significant role in it. Our data provide new insights into the molecular mechanisms regulating stem cell plasticity, by revealing the existence of a “dual-growing” pattern in CRC CSCs.

The spheroid-forming capacity of even a single cell is one of the features of CSCs (Pastrana et al., 2011; Todaro et al., 2007; Vermeulen et al., 2008). In this study, we applied a method that allowed us to precisely track the capacity of not only spheroid formation but also the growth of each SFC in CRC CTOSs with a single-cell resolution: the SSFG assay. The improvement of the *in vitro* culturing conditions (Kondo et al., 2011) compared to those of earlier studies on CSCs from CRC (Todaro et al., 2007; Vermeulen et al., 2008) has contributed to the success of the SSFG assay. The spheroid-forming capacity was varied, but it was generally high in this study, ranging from 27% to 59% (Table S2). Interestingly, the spheroid-forming capacity was similar among the studied clones and subclones, a finding that supports the idea that the variation in spheroid growth is an event that occurs within cell populations that demonstrate spheroid-forming ability.

Recently, the CSC model has been revisited in light of new evidence supporting the status that cancer cells can dynamically fluctuate (Batlle and Clevers, 2017; Hirata et al., 2019; Meacham and Morrison, 2013; Pisco and Huang, 2015). We, herein, demonstrated that the CRC CSCs consist of two distinct but interchangeable subpopulations. The juxtacrine interaction (or cell–cell interaction) with the D-pattern cells as well as the tumor microenvironment was found to be critical in order for the S-pattern cells to acquire the D-pattern phenotype *in vitro*. Our results have also suggested the existence of a gate (Huang, 2009; Pisco and Huang, 2015) that regulates the transition from the S- to the D-pattern in CRC CSCs through a nongenetic process (Figure 7K). In fact, MSI1 seems to be the regulator of this transition.

An overexpression of MSI1 has been reported in different tumor types (Kudinov et al., 2017), including CRCs (Sureban et al., 2008), and MSI1 has been described as a CSC marker in CRC (Forouzanfar et al., 2020; Potten et al., 2003; Schulenburg et al., 2007). We, herein, demonstrated that the expression levels of MSI1 were lower in the S-pattern spheroids, and that these same levels could functionally modulate the transition of the growth status in the studied spheroids (Figure 7). Notably, the spheroid-forming capacity was affected by neither the gene knockout nor the overexpression of MSI1, thus indicating that MSI1 is not essential for the cells to be stem-like. In previous reports, the knockdown of *MSI1* in CRC cells suppressed their capacity of spheroid formation and their tumorigenicity (Sureban et al., 2008; Gao et al., 2014; Li et al., 2015). However, in these cases, the number of spheroids could have been underestimated because of the increased slow-growing populations, and the fact that the MSI1-downregulated cells in these studies did not actually lose their ability to form tumors, but they rather exhibited growth retardation. The molecular mechanisms by which the phenotypically different CSC subpopulations are generated in the D-pattern cells remain to be elucidated. One possible mechanism is that of asymmetric division (Bu et al., 2013; O’Brien et al., 2012; Srinivasan et al., 2016a; Srinivasan et al., 2016b), as MSI1 was originally reported to play a role in asymmetric division (Nakamura et al., 1994).

The majority of current cancer therapies have been designed and developed against fast-growing cancer cells, even when CSCs are targeted (Marine et al., 2020; Recasens and Munoz, 2019). This is despite the fact that resistance to these anticancer therapies has been repeatedly linked to the presence of quiescent or slow-growing CSCs (Boumahdi and de Sauvage, 2020; Cojoc et al., 2015). Given that CSCs fluctuate between different growth states, therapeutic anticancer strategies targeting CSCs should be seriously revisited (Batlle and Clevers, 2017; de Sousa e Melo et al., 2017; Kurtova et al., 2015; Meacham and Morrison, 2013; Rehman et al., 2021; Roesch et al., 2010; Pisco and Huang, 2015). In this study, we revealed that the slow-growing cells play an important role in drug resistance, and that the two identified subpopulations are interchangeable. Our finding that MEK1/2 inhibition cannot affect the growth of the slow-growing cells is in line with a recent work describing a potential side effect of an MEKi as an inducer of stem cell plasticity (Zhan et al., 2019).

The concept of the “drug-tolerant persister” (DTP) has recently emerged as an important driver of therapy failure and tumor relapse. Cancer cells may enter into a nongenetic and reversible DPT state in an attempt to evade cellular death from conventional chemotherapies or molecular-targeted therapies (Guler et al., 2017; Hangauer et al., 2017; Liau et al., 2017; Oren et al., 2021; Sharma et al., 2010). A DTP is a cancer cell that is characterized as being quiescent or slow-growing (Rehman et al., 2021). We observed DTPs in CRC CTOSs treated with high concentrations of chemotherapeutic drugs, and they share similar features with the herein studied slow-growing cells. In fact, the isolated slow-growing cells could be a novel platform for investigating DTPs.

In conclusion, the isolation and stable culturing of the drug-resistant slow-growing CSCs from tumor organoids (without the use of dye retention settings or a combination of markers) can provide a new *in vitro* platform that will allow us to identify and phenotypically characterize drug-resistant slow-growing CSCs. Such a platform could form the basis for the development of new and effective anticancer drugs.

## Methods

### Preparation and culture of cancer-tissue originated spheroid (CTOS)

The study was approved by the Institutional Ethics Committees at Osaka International Cancer Institute (1803125402) and Kyoto University (R1575, R1671). Fresh surgical samples from CRC patients were obtained with the patients’ informed consent. CRC CTOSs were prepared from patient tumor samples or xenografts as previously described with slight modifications (Kondo et al., 2019; Kondo et al., 2011). Briefly, tumors were mechanically minced and incubated for 30-45 minutes in DMEM/Ham’s F12 medium (Fujifilm, 042-30555) containing Liberase DH (Roche, 5401089001) at a final concentration of 0.26 U/mL at 37°C with continuous stirring. DNase I (Roche, 11284932001) was added at 10 μg/mL, followed by an additional 15-minute incubation. The digested solution was serially strained using mesh filters (Falcon® Cell Strainers). Tumor fragments of the 250-500, 100-250, and 40-100 μm fractions were recovered and cultured for 24h in the CTOS medium: DMEM/F-12 with 1x GlutaMAX, 1x StemPro hESC, 1.8% BSA (StemPro™ hESC SFM, Thermo Fisher Scientific, A1000701), and 100 U/mL P-S (Thermo Fisher Scientific, 15140122) in non-treated plates (IWAKI, 1810-006). The next day, CTOSs were washed with HBSS (Fujifilm, 084-08965) to remove cellular debris and cultured either in suspension or embedded in Matrigel GFR (Corning, 354230) in the CTOS medium. The CTOSs prepared from freshly harvested primary CRC tumors (KUCs) were cultured and expanded for 14-21 days in the CTOS medium and subjected to the experiments. For *in vitro* passages, CTOSs were dissociated once a week by the syringe disruption method as previously described (Piulats et al., 2018). Briefly, CTOSs or spheroids were disrupted into smaller fragments by passing them through a 1 mL syringe with a 27 G needle (Terumo, SS-10M2713) at a high flow rate (∼30 mL/min). CTOSs were spontaneously re-formed from these fragments. All experiments were performed at least one day after passaging to avoid the influence of the disruption and remodeling of the spheroids. Within one month of culture, CTOSs were freeze-stocked with StemCell Keep (BioVerde, BVD-VPL-AI-20). ‘CTOS lines’ were defined by the following criteria: 1) growing continuously in culture, 2) generating xenograft tumor (at least 2 passages *in vivo*), and 3) being sufficiently freeze stocked in order to reproduce the experiments. Table S1 presents clinical details regarding the 14 CTOS lines in the CRC panel.

### Xenotransplantation of spheroids

The animal studies were approved by the Institutional Animal Care and Use Committee of Osaka International Cancer Institute (16062411, 17052610, 18060708) and Kyoto University (18564). They were performed in compliance with the institutional guidelines. To generate xenograft tumors of CTOS lines, 2×10^3^ CTOSs were suspended in a 1:1 mixture of medium and Matrigel (Corning, 354234) and subcutaneously injected into the flank of NOD/Scid mice (4-5 weeks old) (CLEA Japan). When tumor volume reached ∼1 cm^3^, mice were sacrificed. For the *in vivo* tumor growth assay, 1×10^3^ CTOSs with similar size (diameter: 40-70 µm) and shape were used for inoculation. Tumor growth was monitored every 2-3 days. Tumor volume was calculated using the following formula: 0.5 x width^2^ x length.

### Single-cell-derived sphere-forming and growth (SSFG) assay

A hundred to a thousand CTOSs were collected and dissociated into single cells by treating with 0.25% Trypsin-EDTA (Thermo Fisher Scientific, 25200072) and DNase I (10 μg/mL) for 10 minutes at 37°C and 1 minute at room temperature, respectively. Then, the cell suspension was gently pipetted a hundred times to promote cell dissociation and filtered through a 35 μm cell strainer (Corning, 352235) to remove cell clusters. The dissociated single cells were diluted in the SSFG medium: CTOS medium containing 2% Matrigel GFR and 10 μM of a ROCK inhibitor, Y-27632 (Selleckchem, S1049), and seeded in a non-treated 384-well plate (Sumitomo Bakelite, MS-9384U) with a ratio of 1 cell per well (50 μL/well) using an E1-ClipTip electronic pipette (Thermo Fisher Scientific, 4672060BT). Within two hours after cell seeding (day 0), the presence of one cell per well was confirmed by image acquisition using the LEICA DMI4000B microscope (Leica Microsystems) and Lumina Vision software (Mitani Corporation). Single cells with an area greater than 300 μm^2^ on day 0 and wells containing multiple cells were excluded from the subsequent analyses. The growth of single cells into spheroids was monitored by image acquisition. Fresh SSFG medium without Y-27632 (30 μL/well) was added on day 7. The ability to form spheroids as well as the growth were evaluated on day 13 for C45 clones and on day 20 for other lines unless otherwise noticed, by measuring the surface area of each single-cell-derived spheroid using the acquired pictures and ImageJ Fiji software (https://imagej.net/software/fiji). Spheroid-forming capacity was calculated and expressed as the percentage of single cells able to grow and form spheroids. Mean ± SD from at least three independent experiments is shown. For the experiments with chimeric spheroids, the single cell-derived spheroids with different fluorescence were assessed by image acquisition using the LEICA DMi8 microscope and LAS X software (Leica Microsystems). For the experiments with a MEK1/2 inhibitor (PD0325901, Selleckchem, S1036), dispersed single cells were diluted in the SSFG medium containing 10 or 100 nM of the inhibitor. The SSFG medium supplemented with 0.1% of DMSO (Millipore Sigma, D5879) was used as a control. Fresh SSFG medium with the same doses of DMSO or MEK1/2 inhibitor was added after 7 days of culture.

### Isolation and culture of the slow- and dual-growing spheroids

At the first round of the SSFG assay, the selected single-cell-derived slow- and fast-growing spheroids were picked up and individually cultured and expanded *in vitro* in the CTOS medium. A subsequent round of SSFG assay was performed for each selected tumor spheroid clone to evaluate its growing pattern: slow- or dual-growing phenotype. For the additional rounds of the SSFG assay, the assay was sequentially performed for each clone and indicated as round one (x1), two (x2), and three (x3). Between the rounds, a pool of spheroids with similar size was collected and subjected to the next round of the SSFG assay.

### Spheroid co-culture system

To generate chimeric spheroids, fluorescent-labeled spheroids were dissociated into single cells and mixed at a 2:1 ratio (1×10^4^ cells of EGFP-labeled C45-4SR : 5×10^3^ cells of mCherry-labeled C45-4D or 1×10^4^ cells of tRFP-labeled sgRNA MSI1-C45-4D : 5×10^3^ cells of wild-type C45-4D). The cell mixture was suspended in the SSFG medium and dispensed into the round-bottom non-treated 96-well plate (Greiner bio-one, 650185). The plate was centrifuged at 400x g for 3 min to facilitate aggregation and incubated for 7 days. The chimeric spheroids were collected and subjected to the SSFG assay. For the DAPT treatment experiments, the fluorescent-labeled spheroids were pretreated overnight with 50 µM of DAPT (Abcam, ab120633) or 0.1% of DMSO as control. Then, the chimeric spheroids were generated in the presence of DAPT (50 µM) or DMSO (0.1%), cultured for 7 days, and subjected to the SSFG assay. For the co-culture system without physical cell-cell interaction, 1×10^3^ cells for each clone were separately embedded in 7 µL of Matrigel GFR, solidified as a droplet in a non-treated 24-well (IWAKI, 1820-024), overlaid with CTOS medium containing 10 μM Y-27632, and cultured for 7 days. The EGFP-labeled-C45-4SR spheroids were collected and subjected to the SSFG assay. For control experiments, the same number of EGFP-labeled C45-4SR cells were cultured alone in the same way.

### CTOS drug sensitivity assay

The CTOS drug sensitivity assay was performed as previously described (Kondo et al., 2019) with slight modifications. Briefly, spheroids with similar size and shape (diameter: 40-100µm) were collected and seeded in non-treated 24-well plates at a density of 1×10^2^ per well. Spheroids were cultured for one week in the CTOS medium containing the indicated dose of the drugs or 0.1% of DMSO as control (n = 3 wells for each condition). Pictures of the entire well were captured on day 0 and day 7. The viabilities of the spheroids were evaluated using the CellTiter-Glo assay (Promega, G7570). ATP content was measured at day 7 and adjusted to the control group. The replicas of the spheroids at day 7 were subjected to the SSFG assay. The 5-FU (Kyowa Kirin Co., Ltd) and gemcitabine (Eli Lilly Japan) compounds were provided by the Department of Pharmacy, Osaka International Cancer Institute.

### Vector construction and gene transfer

The following lentivirus-expressing vectors were used: 7TFC-mCherry (Addgene #24307, a gift from Roel Nusse); pLX304 (Addgene #25890, a gift from David Root); and pL-CRISPR.SFFV.tRFP (Addgene #57826, a gift from Benjamin Ebert). For generating the pLX304-EGFP vector, the EGFP sequence was transferred from the pDONR221_EGFP plasmid (Addgene #25899, a gift from David Root) into the pLX304 plasmid by the Gateway cloning system (Thermo Fisher Scientific, 11791020). For MSI1 overexpression, human MSI1 cDNA was amplified with the primers of MSI1_BamHI-F and MSI1_stopdead_XbaI-R (Table S6) using the PrimeScriptTM 1st strand cDNA Synthesis (Takara, 6210A) and PrimeSTAR ® Max DNA Polymerase kits according to manufacturer instructions (Takara, R045A). The PCR product was cloned into the pENTR4 no ccDB plasmid (Addgene #17424, a gift from Eric Campeau & Paul Kaufman) and transferred into the pLX304 plasmid by the Gateway cloning system. The lentiviral vectors were transfected (Roche, 6366236001; Addgene #12259 and #12260, both gifts from Didier Trono) as previously reported (Onuma et al., 2021). For the pLX304-GFP and -MSI1, blasticidin (2 µg/mL) (Millipore Sigma, 15205) selection was performed. For the pLX304-EGFP and 7TFC-mCherry, the green (EGFP)- and red (mCherry)-positive cells from the reconstituted spheroids, respectively, were sorted and expanded. For *MSI1* knockout, single-guide RNA (sgRNA) or a non-targeting (sgCtrl) sgRNA was cloned into a pL-CRISPR.SFFV.tRFP expression vector (Addgene #57826, a gift from Benjamin Ebert) as described by Heckl et al. (Heckl et al., 2014). The sgRNA sequences were designed using the Broad Institute sgRNA designer tool (http://portals.broadinstitute.org/gpp/public/analysis-tools/sgrna-design). The sequences of the sgRNA oligos are shown in Table S6. Single cells derived from C45-4D spheroids were transduced with the lentiviral Cas9-expression vector (pL-CRISPR.SFFV.tRFP) containing the MSI1 sgRNA or the non-targeting sgRNA as control. The red (tRFP)-positive cells from the reconstituted spheroids were sorted and expanded.

### RNA isolation and semi-quantitative real-time PCR

Spheroids were cultured for 7 days from single cells embedded in Matrigel GFR (5×10^2^ cells / 7 µL of Matrigel GFR) in CTOS medium containing 10 μM of Y-27632. Total RNA was extracted with the RNeasy mini kit (Qiagen, 74106) plus on-column DNAse I digestion (Qiagen, 79254). Semi-quantitative real-time PCR was carried out (Thermo Fisher Scientific, 18080085 and 4385617) as previously described (Endo et al., 2017). Data are presented as the mean ± SD of three replicates. The primer sequences are presented in Table S6.

### Microarray and GSEA analysis

Spheroids were cultured for 7 days from single cells embedded in Matrigel GFR (5×10^2^ cells / 7 µL of Matrigel GFR) in CTOS medium containing 10 μM of Y-27632. Total RNA was extracted as previously described. Microarray analysis, from three biological replicates, was performed with the GeneChip Human Gene 2.0 ST Array. Signals were quantified and normalized with the RMA algorithm. For gene set enrichment analysis (GSEA), software was downloaded from the Gene Set Enrichment Analysis website (http://www.broad.mit.edu/gsea/downloads.jsp). GSEA was performed using the c-Myc target (Zeller et al., 2003) gene set to identify enriched/depleted signatures. Gene sets with an FDR< 0.25 and a nominal P-value of <0.05 were considered significant. Volcano plots were generated with the GraphPad Prism version 9 (GraphPad Software) using the transcriptome data and setting the threshold for the log_2_(Fold Change) as ± 0.59 (Fold change < 0.67 = -0.59; Fold change > 1.5 = +0.59) and − Log _10_(p-value) as 1.3 (>1.3 = P < 0.05). To generate a heatmap, gene expression data of day 8 samples were analyzed. After omitting the probes without gene name, the probes with average expression level at top 40% were extracted. Then, FDR was calculated for 6 slow samples vs 3 fast samples. Probes with FDR<0.1 were extracted to generate heatmap (heatmap3: R version 3.6.3 (http://www.R-project.org)).

### Western blot analysis

Spheroids in suspension or single cell-derived spheroids embedded in Matrigel GFR (5×10^2^ cells in 7 µL of Matrigel GFR) were cultured for 7 days in the CTOS medium. Western blotting analyses were performed as previously described (Endo et al., 2017). The total AKT (#2920), pAKT-S473 (#9271), MEK1/2 (#9122), pMEK1/2-S217/221 (#9121), ERK1/2 (#9107), pERK1/2-T202/Y204 (#4370), GAPDH (#2118), S6 (#2212), p-S6 (#2211), c-Myc (#5605), and NICD (#4147) antibodies were obtained from Cell Signaling Technology. MSI1 (ab52865) antibody was obtained from Abcam. Secondary anti-rabbit (#7074) and -mouse (#7076) antibodies were obtained from Cell Signaling Technology. All antibodies were used at the producer’s suggested concentrations.

### Quantification and statistical analysis

Statistical analyses were carried out with GraphPad Prism 9 (GraphPad Software). Significance was tested with the unpaired Student’s t-test for single comparisons, and with one-way or two-way ANOVA analysis followed by Tukey’s or Bonferroni’s test for multiple comparisons as indicated. For the analyses of SSFG assay results that did not show a normal distribution, a non-parametric test, the Mann-Whitney test, was used. P-values < 0.05 were considered significant. Violin plots showing the area of growing spheroids and frequency distribution analysis of their size (indicated as a percentage) were performed using the GraphPad Prism version 9. Representative data of three independent experiments are shown for the violin plots. Bimodal distribution was tested by Silverman’s bootstrap test with the null hypothesis that the kernel density has one mode, *p < 0.1, using VISUAL-SILVERMAN (Kusuhashi and Okamoto, 2015). Ns, not statistically significant; *, P < 0.05; **, P < 0.01; ***, P < 0.001; and ****, P < 0.0001.

## Data availability

Further information and requests for resources and reagents should be directed to and will be fulfilled by the lead contact, Masahiro Inoue. There are restrictions to the availability of CRC CTOS lines through Material Transfer Agreement requirements at Kyoto University specific to the CRC CTOS and the subclones. The accession number for the gene expression microarray data reported in this study is GEO: GSE185012.

## Acknowledgments

This work was supported by a Grant-in-Aid from P-CREATE, a Japan Agency for Medical Research and Development, Japan, 20cm0106203h0005 (M.I., R.C., J.K., K.O.); by the Cabinet Office, Government of Japan, Public/Private R&D Investment Strategic Expansion Program (PRISM) (Y.T., M.K.); and by the RIKEN Junior Research Associate Program (Y.T.). We thank Atsuko Takagi and Kasumi Ota for their technical assistance.

## Author contributions

R.C. and M.I. designed the experiments and wrote the manuscript. R.C., K.O., M.K., Y.T., and K.K. conducted the experiments. J.K. interpreted the data obtained. K.I. and M.O. performed the statistical analyses. K.O. and M.I. supervised the project.

## Competing interests

J.K. and M.I. belong to the Department of Clinical Bio-resource Research and Development at Kyoto University, which is sponsored by KBBM, Inc. M.I. is an inventor of the patents related to the CTOS method.

## Supplementary information

**Supplementary Figure S1.**
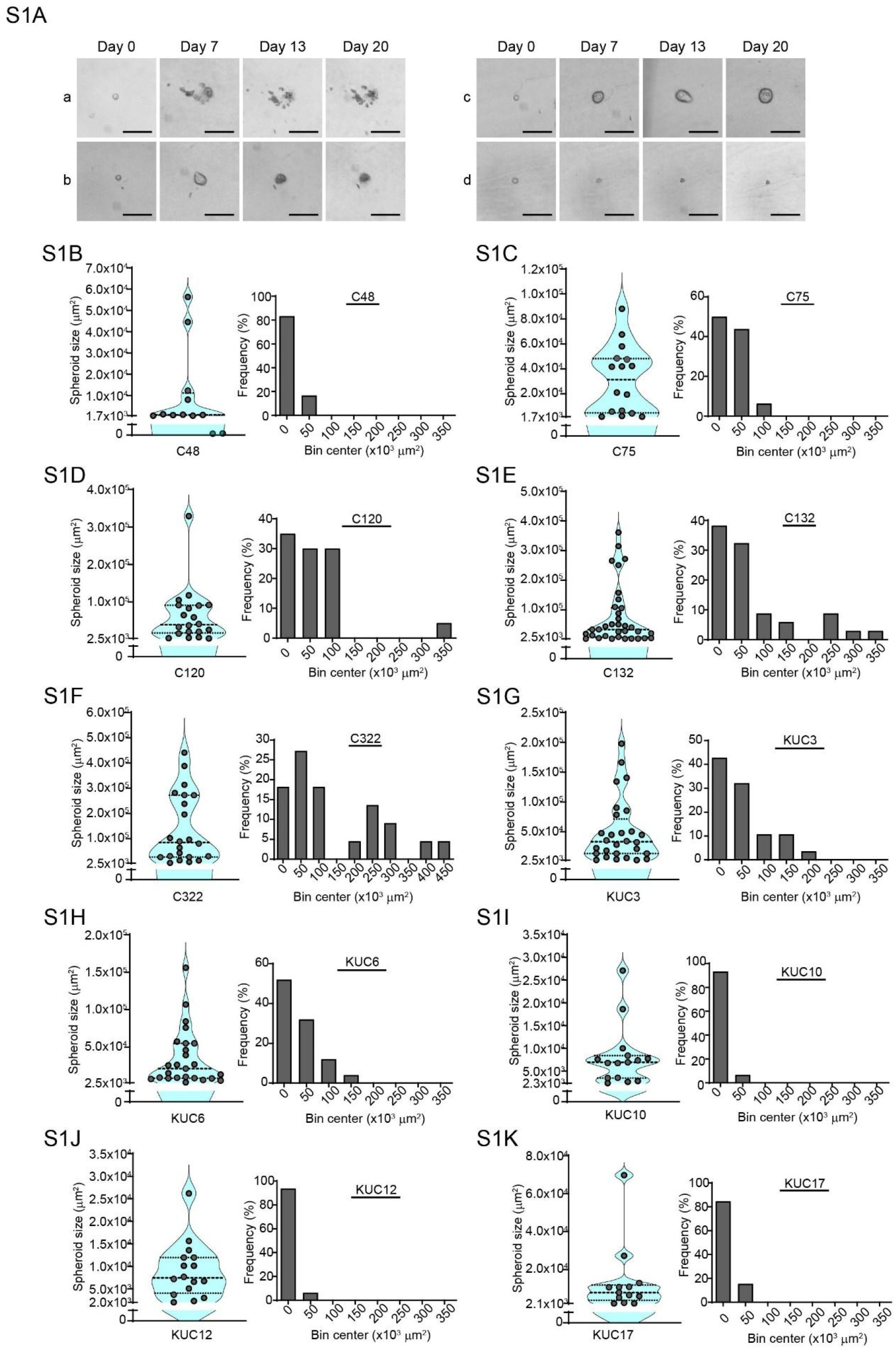
Images of non-growing spheroids and the SSFG assay results of additional CRC organoids. (**S1A**) Representative phase-contrast images of non-growing spheroids from Figure 1C at the indicated days of culture. Scale bars, 100 μm. (**S1B-S1L**) Violin plots and frequency distribution analysis of the SSFG assay in the indicated CRC CTOS lines.

**Supplementary Figure S2.**
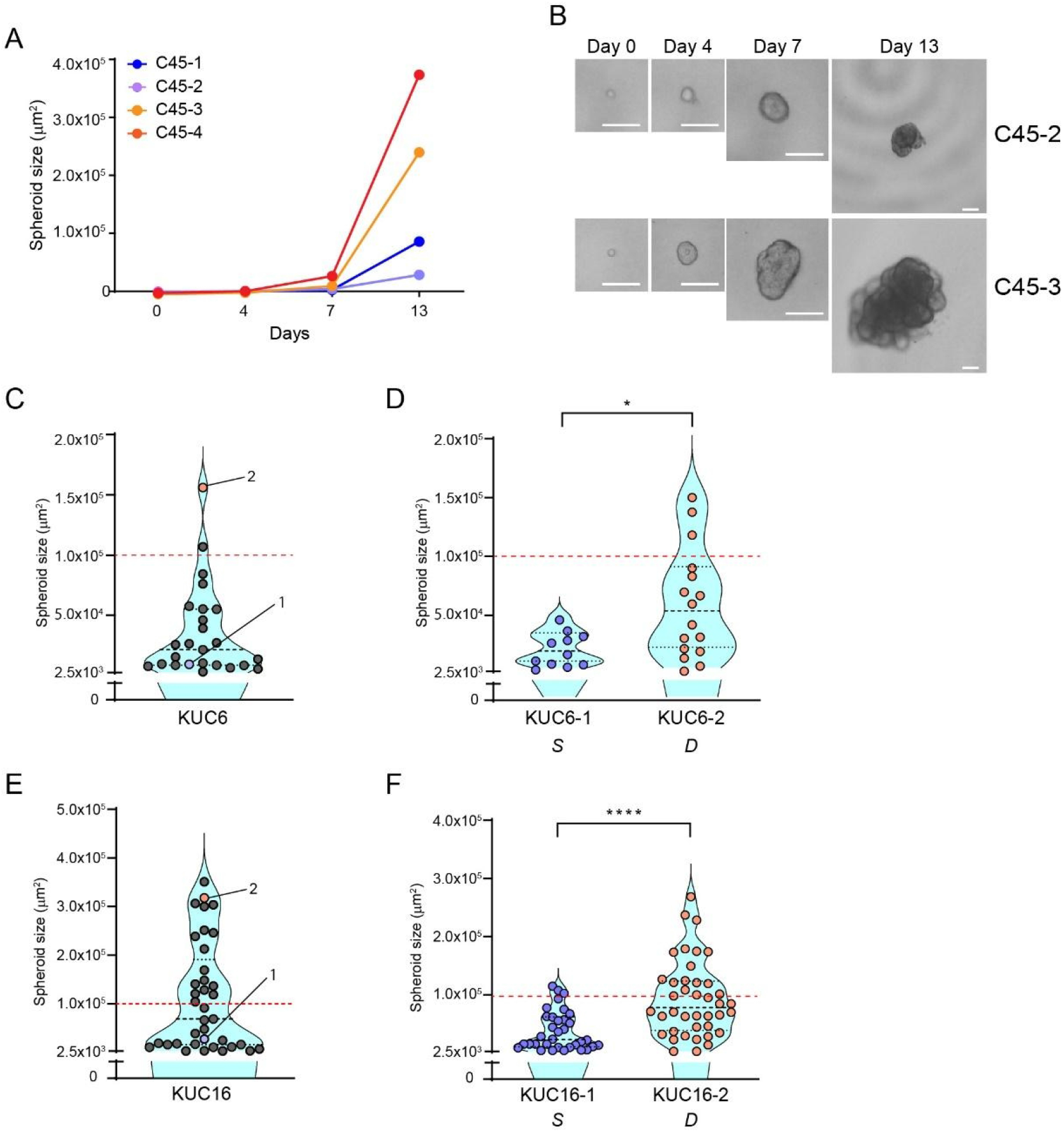
CRC spheroids exhibit different growth capacities with preserved growth potential. (**S2A**) Growth curve of the four selected spheroids in Figure 2A. The area of spheroids at different days of culture is indicated. (**S2B**) Time-course images of C45-2 and -3. Scale bars, 100 μm. (**S2C-S2F**) Violin plots of the SSFG assay for primary CTOSs and the selected clones for KUC6 (S2C, S2D) and KUC16 (S2E, S2F) lines. The selected clones, KUC6-1 and -2, and KUC16-1 and -2, are indicated in S2C and S2E, respectively. *S*, S-pattern; *D*, D-pattern.

**Supplementary Figure S3.**
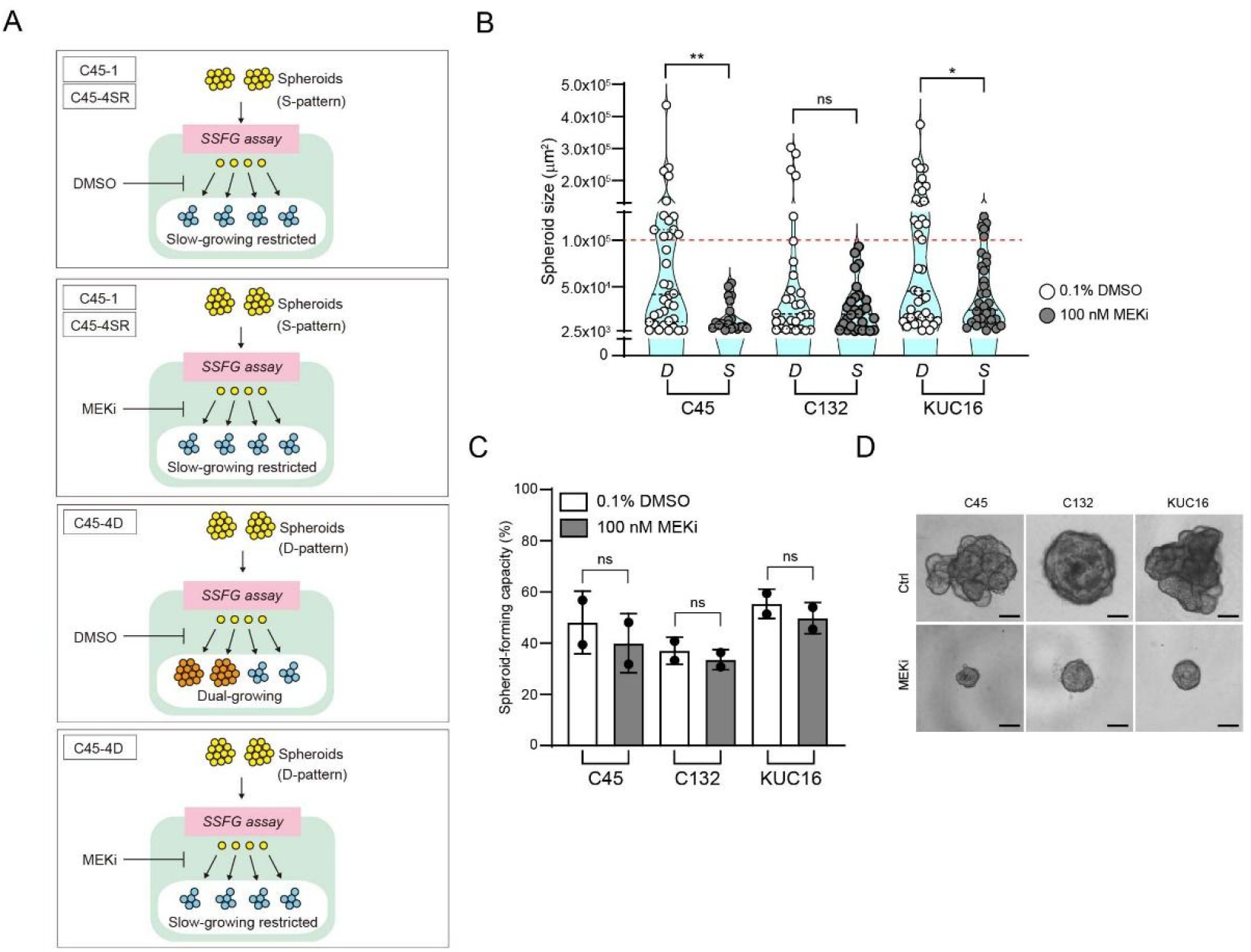
Slow-growing CRC spheroids show resistance to a MEK1/2 inhibitor with a single-cell resolution. (**S3A**) Schematic overview of the experimental design and the results of Figure 5B-5F. (**S3B-S3D**) Violin plot of the SSFG assay (S3B), spheroid-forming capacity (S3C), and representative images (S3D) for the indicated CRC CTOS lines comparing DMSO (0.1%)-treated with MEKi (100 nM)-treated for 13 days. C45 represents parental C45. Scale bars, 100 μm (S3D). *S*, S-pattern; *D*, D-pattern.

**Supplementary Table S1.**
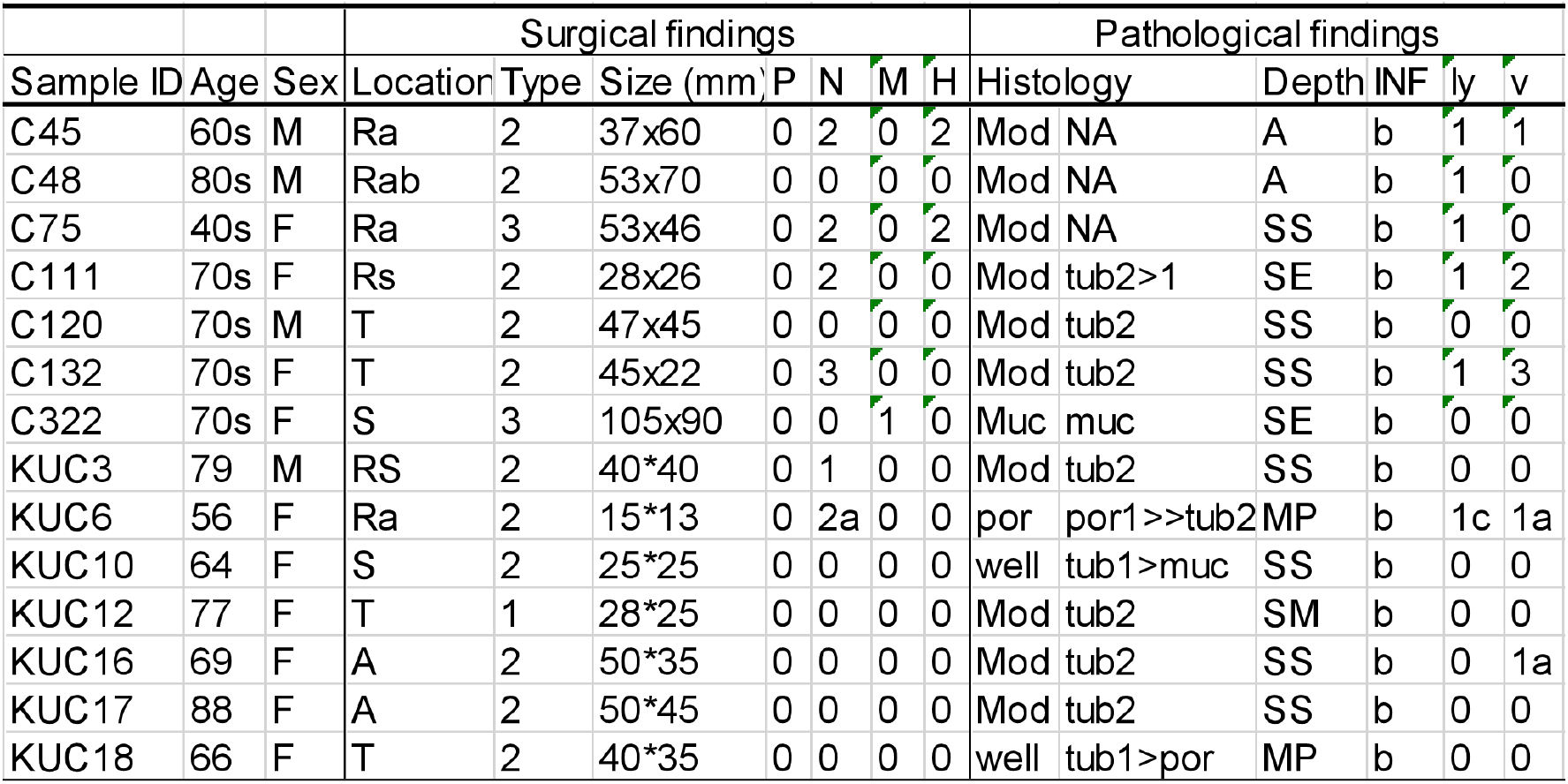
Clinical information of the CTOS lines and the primary tumors.

**Supplementary Table S2.** Summary of SSFG assay results in 14 CRC CTOS lines. C45 was also evaluated after 20 days of culture. Spheroid-forming capacity, minimum and maximum growth size, and growth range of growing spheroids are indicated.

| Table S2: SFC and growth range |  |  |  |  |
| --- | --- | --- | --- | --- |
| CTOS line | SFC (%) | Minimum growth size (x10<sup>3</sup> μm<sup>2</sup>) | Maximum growth size (x10<sup>3</sup> μm<sup>2</sup>) | Growth range (Fold change) |
| C45 | 59 | 2.7 | 616 | 228 |
| C48 | 19 | 1.7 | 56 | 33 |
| C75 | 20 | 1.7 | 88 | 52 |
| C111 | 40 | 2.6 | 47 | 18 |
| C120 | 47 | 2.6 | 329 | 128 |
| C132 | 51 | 2.6 | 583 | 228 |
| C322 | 49 | 3.7 | 440 | 120 |
| KUC3 | 36 | 2.7 | 198 | 74 |
| KUC6 | 36 | 3.2 | 156 | 49 |
| KUC10 | 27 | 2.3 | 23 | 12 |
| KUC12 | 30 | 2 | 26 | 13 |
| KUC16 | 59 | 2.6 | 375 | 142 |
| KUC17 | 38 | 2.1 | 70 | 33 |
| KUC18 | 33 | 2.8 | 98 | 34 |
| SFC: Spheroid forming capacity |  |  |  |  |

**Supplementary Table S3.**
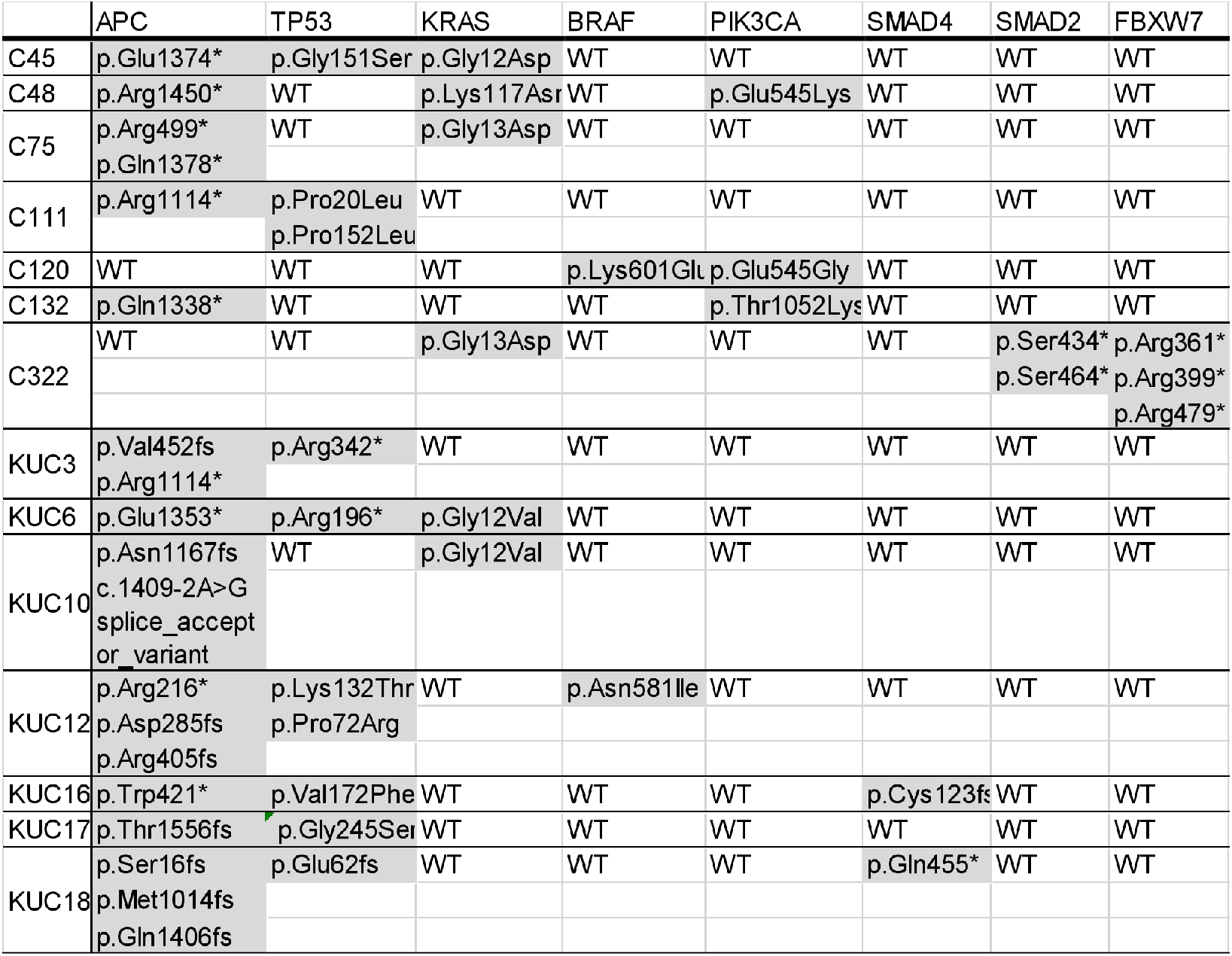
Mutational status of frequently mutated genes in CRC for the CTOS lines and primary tumors. Mutations are highlighted in gray.

**Supplementary Table S4.**

List of differentially-expressed genes between C45-4D and C45-1, C45-4D and C45-4SR, C45-4SR and C45-1, in Figures 6A-6D. Upregulated genes (fold change < 0.67, p<0.05) and downregulated genes (fold change > 1.5, P<0.05) are shown in each case.

**Supplementary Table S5.**

List of differentially-expressed genes in C45-4D commonly compared to C45-1 and C45-4SR in Figure 6H.

**Supplementary Table S6.**
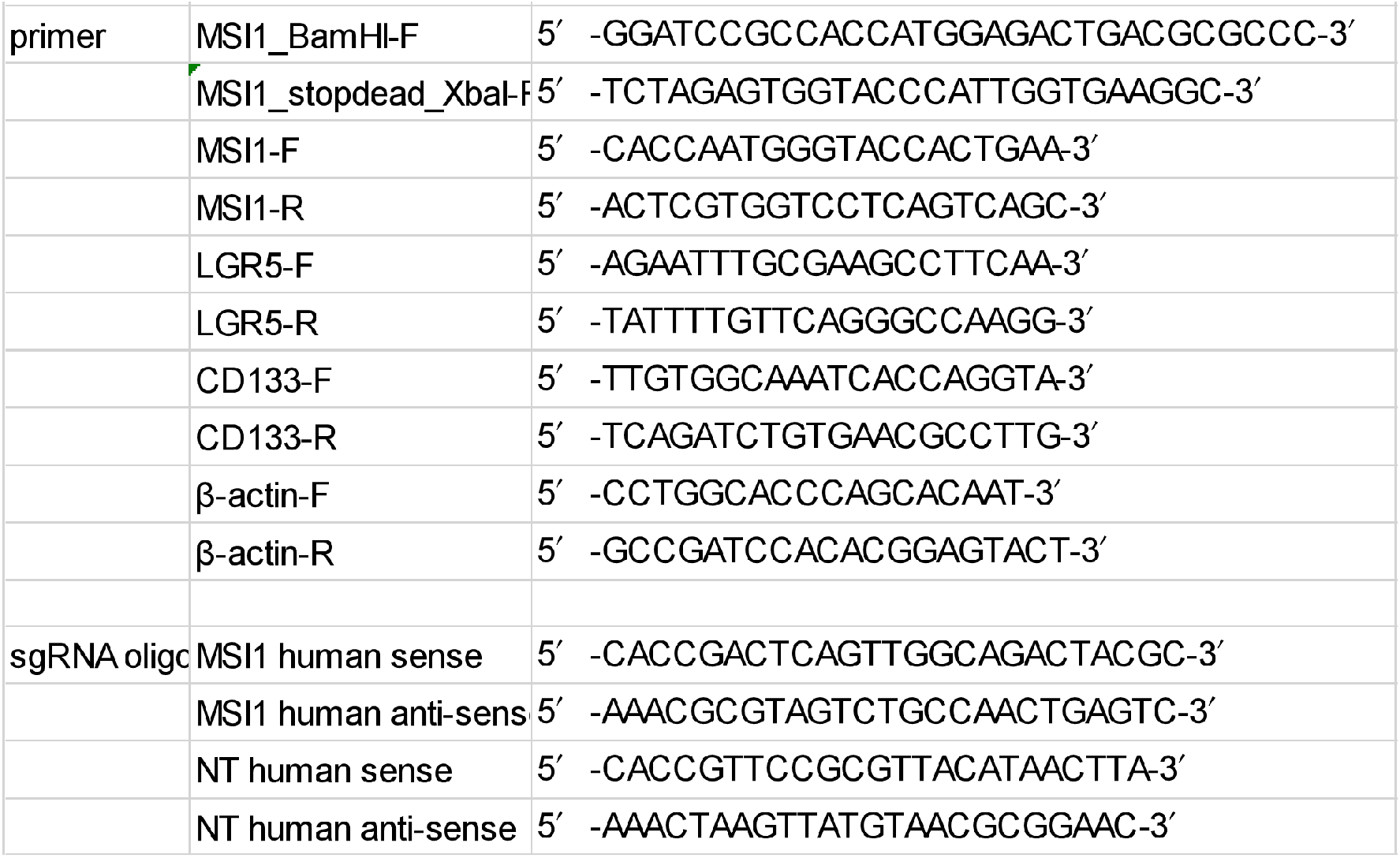
Sequences of the PCR primers and sgRNA oligos.

